# Calibrating Classifiers Across Covariates: Hierarchical Vocal Density Estimation for Acoustic Monitoring at Scale

**DOI:** 10.64898/2026.09.08.747031

**Authors:** Louis Freeland-Haynes, Matthew J. Weldy, Amanda K. Navine, Patrick J. Hart, Tom Denton, Justin Kitzes

**Author notes:** These authors contributed equally and share senior authorship (Tom Denton, Justin Kitzes).

## Abstract

1. Passive acoustic sensors and machine learning classifiers offer a scalable approach to monitoring wildlife populations. These classifiers typically detect whether a species is present in a short time-window of audio; ecologists, however, want ecologically meaningful measures such as indices of site-level abundance. Vocal density, the proportion of time-windows containing a vocalization, is a useful abundance proxy. Recovering vocal density from classifiers requires calibrating their outputs against expert-labeled data. Because classifier performance varies between sites, a single global calibration yields overconfident site-level estimates, yet calibrating every site independently requires prohibitive labeling effort. Estimating site-level vocal density at the scale of modern sensor networks demands a label-efficient alternative.
2. We present a Bayesian hierarchical extension of Platt scaling, a commonly used calibration method, that calibrates classifiers at the site level while borrowing strength across sites. Where site-level covariates are available, a Gaussian-process prior lets the model learn how calibration varies across covariate space. Averaging calibrated per-clip probabilities across a site’s recordings yields a posterior distribution over vocal density, and we show how to carry that uncertainty into downstream regressions of vocal density on environmental covariates.
3. Across simulations and two fully annotated field datasets from Hawai’i and the Pacific Northwest, our model attained desired credible-interval coverage of vocal density estimates with narrower intervals than independent site-level calibration at equal labeling effort, and lower mean squared error. Global calibration, by contrast, failed to attain coverage except when simulated sites were genuinely homogeneous. Unlike site-level calibration, our model enables calibration at sites with zero labeled data, and incorporating covariates further improved precision when site heterogeneity was covariate-driven. Applied to a 283-site dataset in Pennsylvania with only two labeled clips per site, we recovered known habitat associations for the declining Wood Thrush (*Hylocichla mustelina*).
4. Our method provides a principled, label-efficient route from bioacoustic classifier scores to site-level abundance indices with well-quantified uncertainty, making rigorous ecological inference feasible at the scale at which acoustic sensor networks are now deployed.

## 1 Introduction

Monitoring wildlife populations and understanding what drives changes in their occurrence and abundance are central problems in ecology and conservation. Sensors such as camera traps and acoustic recorders are able to generate vast amounts of data, and pairing them with machine-learning classifiers for automated detection offers an appealing path toward automated monitoring at large spatial and temporal scales. Bioacoustics is one field where this approach has been adopted especially rapidly: acoustic datasets are accumulating globally (Darras et al., 2025; Sugai et al., 2019), freely available classifiers now enable automated recognition of bird species worldwide (Kahl et al., 2021; van Merriënboer et al., 2025), and other taxa are increasingly represented (Mac Aodha et al., 2022; Hagiwara et al., 2022).

A central challenge is how to estimate ecologically useful measurements from the outputs of these machine learning classifiers. Currently widely used automated classifiers are designed to detect whether a species of interest is present in an individual data sample. In bioacoustics, that typically means predicting whether a short audio time-window (often 3-5 seconds) contains a vocalization of interest (Stowell, 2022). Importantly, the scale of these individual predictions does not match common scales of ecological interest, such as point-level occupancy or abundance.

Abundance, the count of individuals present at a location in a defined time window, is often the most important variable of interest in most population-level ecological research (Nichols and MacKenzie, 2004). Ecologists typically wish to estimate abundance at the point or site-level, which in the context of bioacoustics may correspond to a single autonomous recording unit (ARU) or a set of ARUs covering an area of interest. Directly estimating abundance from acoustics is not trivial (Pérez-Granados and Traba, 2021), however vocal density - the proportion of time-windows containing a vocalization -has been shown to be a useful proxy for abundance for some species (Pérez-Granados et al., 2019; Jarrett and Willis, 2025; Navine et al., 2024a).

Estimating site-level vocal density from the clip-level outputs of a classifier is complicated by the properties of imperfect classifiers. Classifiers produce continuous scores between 0 and 1 that reflect the model’s degree of belief that a class of interest is present in a sample. The most straightforward estimator of vocal density is to apply a decision threshold, treating samples above that threshold score as detections and those below as non-detections, and take the proportion of detections as vocal density (Knight et al., 2017). For any imperfect classifier, however, the confidence score distributions of true positives and true negatives overlap and no threshold cleanly separates them: any choice of threshold trades false positives against false negatives, and false positive and false negative rates will vary by site (Metcalf et al., 2022). False positive abundance models have been developed to handle such data (Udell et al., 2024; Clement et al., 2022; Doser et al., 2021), however the approach of thresholding necessarily collapses each clip to a single above-or below-threshold classification, discarding information carried in the continuous scores (Kitzes et al., 2026).

An alternative to threshold-based methods is to interpret the continuous scores probabilistically. If the outputs of a classifier are marginally calibrated then thresholding is unnecessary: the proportion that are positive can be obtained directly by averaging the scores across all samples (Moreo, 2025). In reality, the outputs of convolutional neural networks (CNNs) are typically poorly calibrated (Guo et al., 2017), and this has been demonstrated for bioacoustic classifiers (Schwinger et al., 2026). Turning classifier outputs into usable probabilities therefore requires an additional step of post-hoc calibration. How best to calibrate a classifier is an active area of research (Silva Filho et al., 2023). Post-hoc calibration methods use a labeled subset of data (the calibration set) to learn a mapping from raw score to the probability that an audio window contains the target species. Several approaches have been applied in ecological settings, including Platt scaling (Platt, 1999; Wood and Kahl, 2024), temperature scaling (Dussert et al., 2025), and logarithmic histogram binning (Navine et al., 2024b).

All calibration, however, requires labeling effort. A calibration set of data must be labeled by an expert, and in practice expert annotation time is often the limiting constraint. Approaches that achieve reliable calibration from fewer labels are therefore especially valuable. Navine et al. (2024b) made notable progress on this problem with logarithmic histogram-binning. They bin the observed score distribution on a logarithmic scale and estimate positive prevalence within each bin. This stratified sampling heuristic directs the bulk of labeling effort toward the highest-scoring clips while ensuring coverage across the score range -an approach well-suited to bioacoustics, where positives are rare. In contrast, uniform sampling when positives are rare can result in all or almost all annotated clips being negatives, yielding little information about the score-to-probability mapping in the range that matters.

Log-binning reduces the labels needed per unit (such as a site), but total annotation effort still scales linearly with the number of units. If site-level vocal densities are desired, calibrating each site independently is prohibitive at scale. Modern sensor networks may span hundreds to thousands of sites (Sugai et al., 2019; Darras et al., 2025). If, for example, estimating vocal density at a desired precision required 50 labeled clips, a study investigating 10 species at 1,000 sites would require 500,000 expert-labeled clips. Pooling every site to perform a single global calibration is tempting but ill-advised. Sites differ in recorder quality, background noise, and the presence of co-occurring species, all of which can shift a classifier’s performance (Metcalf et al., 2022; Navine et al., 2024b).

Using a global calibration to infer densities at individual sites risks an ecological fallacy (Freedman, 1999) by assuming that an aggregate-level relationship holds for its constituent units. Existing approaches thus sit at two ends of a trade-off: global calibration is cheap but biased and cannot adequately account for site-level uncertainty, while site-level calibration is accurate but scales poorly. Neither enables estimation of site-level vocal density at the scales of deployment that motivate automated monitoring.

To resolve this trade-off, we propose a covariate-informed calibration method that uses limited labeled data to calibrate a classifier at the site level, producing estimates of site-level vocal density with appropriately quantified uncertainty. The approach extends Platt scaling in a hierarchical Bayesian model, borrowing strength across sites so that each site’s calibration is informed both by its own labeled data and by the broader population of sites. Site-level covariates can be incorporated to explain variation in calibration between sites.

We evaluate our method on three types of data of increasing realism. First, on simulated data where ground truth is known, we verify that our model attains nominal coverage with narrower intervals than independent site-level calibration, effectively reducing the amount of labeling effort needed to reach a desired precision. Second, on two fully annotated field datasets, from Hawai’i and the Pacific Northwest, where exhaustive labeling gives us realized vocal density, we show that these gains hold on real-world model predictions on audio. Finally, we apply the method in its intended setting to a partially annotated dataset of over 200 sites from the eastern deciduous forest, recovering covariate effects that agree with known ecological relationships from only two labeled clips per site.

## 2 Methods

Our aim is to estimate site-level *vocal density*, the proportion of audio clips at a site that contain a class of interest. To do so, we leverage the predictions of a bioacoustic classifier, whose scores are not calibrated probabilities and whose calibration may differ between sites. We do so by extending Platt scaling, a two-parameter logistic calibration (Platt, 1999), into a Bayesian hierarchical model (Figure 1). Each site receives its own intercept and slope, drawn from a shared prior, so that each site’s calibration is informed both by its own labeled data and by the other sites. In the absence of covariates this is a simple random effect. Where site-level covariates are available, we use a Gaussian process to let the shared prior depend zon covariates, learning how calibration varies across covariate space and improving the precision of site-level estimates. Our approach can be summarized in the following steps:

1. Fit the calibration model using the set of labeled clips.
2. Apply each site’s fitted calibration to the unlabeled clips to produce calibrated per-clip probabilities.
3. In a posterior predictive simulation, aggregate predicted labels on the unlabeled clips from the calibrated probabilities (and any known labels from labeled data). This yields a posterior over site-level vocal densities.
4. When the goal is ecological inference, fit a second stage regression of vocal density against environmental covariates.

**Figure 1.**
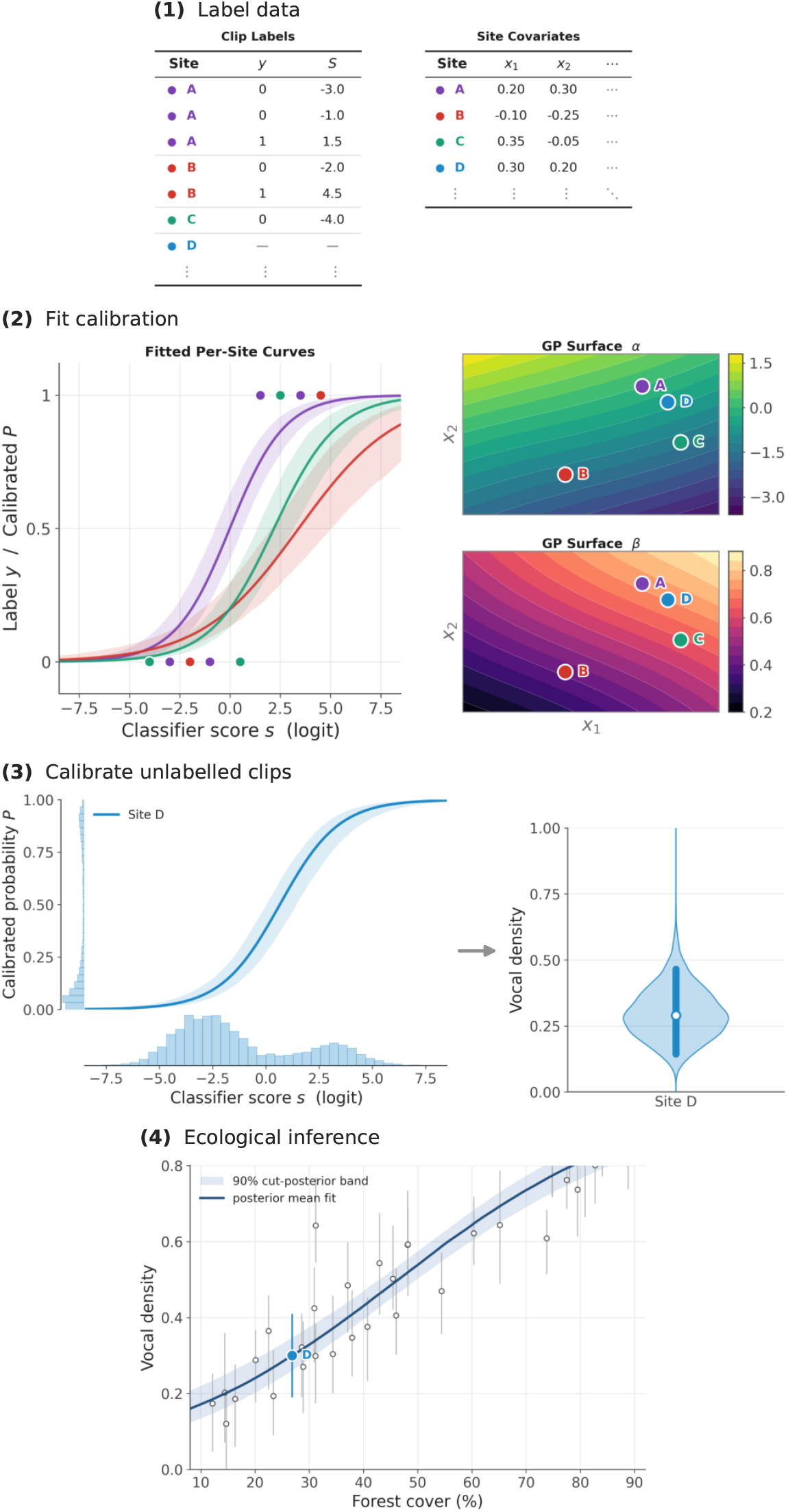
Conceptual overview of our hierarchical calibration method. **(1) Label data**. An expert labels clips (selected efficiently via a labeling strategy). The label *y*, logit-scale score *S*, and site-level covariate values *x*_1_, *x*_2_, … are passed to the calibration model. **(2) Fit calibration**. Labeled data are used to fit per-site logistic curves, using a Bayesian hierarchical model. To incorporate information from site covariates, Gaussian Processes model how a site’s intercept *α* and slope *β* vary across covariate space. **(3) Calibrate unlabeled clips**. Scores on unlabeled clips are pushed through a sites posterior *α* and *β* to produce calibrated probabilities. These imply a posterior over vocal density. This is possible even if a site has no labeled data (as in site D). **(4) Ecological inference**. Per-site posteriors feed into a second-stage regression on an environmental covariate (forest cover), propagating uncertainty. The posterior of vocal density at site D is highlighted.

Because inference is fully Bayesian throughout, each stage yields posterior distributions, which we summarize with highest-posterior-density credible intervals.

### 2.1 The calibration model

Bioacoustics is a multi-label context, as a single audio window may contain several classes. We therefore treat our calibration problem as a collection of binary classifiers, calibrating one target class at a time. The description below applies to a single focal class. For each labeled clip j at site i, let *S*_*ij*_ be the classifier score on the logit scale, and *y*_*ij*_ in *{*0, 1*}* indicate whether the class of interest is present. We model the outcome with a site-specific logistic (Platt) calibration:

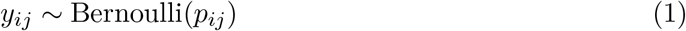

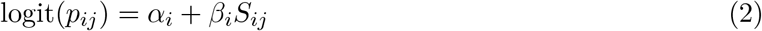

where *α*_*i*_ and *β*_*i*_ are site-level intercept and slope. Global Platt scaling is the special case *α*_*i*_ = *α, β*_*i*_ = *β* for all sites. Independent site-level calibration fits each (*α*_*i*_, *β*_*i*_) separately with no pooling. Our model uses a hierarchical approach: each site has its own slope and intercept, drawn from a shared prior that pools information across sites.

#### Hierarchical prior over site-level calibrations

We first describe a simpler model that can be fit without covariates. Site is treated as a random effect. Each site’s intercept and slope are drawn from a shared distribution, so that estimates are partially pooled:

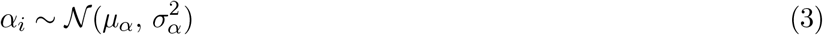

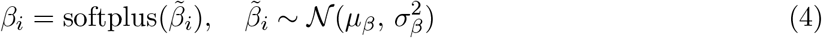

The softplus transform, softplus(*x*) = log(1 + *e*^*x*^) maps the slope to the positive reals, ensuring the calibration function is monotonically increasing in classifier score -i.e. that the classifier is at worst no-skill, and higher scores correspond to higher calibrated probabilities.

We extend this covariate-free model to leverage information in site-level covariates that may be associated with calibration. A confusion sound or species, for instance, may be present in only part of covariate space, shifting calibration there but not elsewhere. Because the ways covariates may affect calibration are varied and not known in advance we use a Gaussian Process (GP); a flexible non-parametric approach to exploit this information and improve precision of site-level estimates.

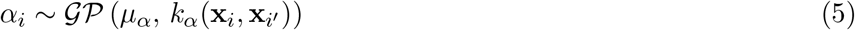

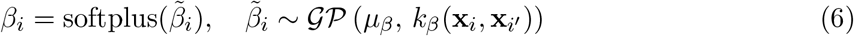

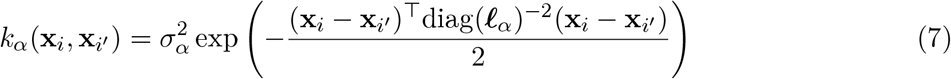

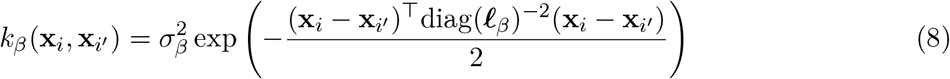

Each GP maps a point in *d*-dimensional covariate space to a distribution over plausible site-level intercepts (or slopes). The covariance functions *k*_*α*_ and *k*_*β*_ are squared-exponential kernels with independent length-scales for each covariate dimension (Wipf and Nagarajan, 2007). This acts as a prior favouring smooth functions of the covariates: sites with similar covariate values are expected to have similar calibrations, and the similarity between two sites decays with a squared-exponential of their distance in covariate space. The per-dimension length-scales ***L***_*α*_, ***L***_*β*_ allow the model to learn which covariates most strongly influence calibration, down-weighting uninformative covariates through large fitted length-scales. To each covariance matrix we add a per-site nugget on the diagonal, which allows sites with identical covariates to differ in their calibrations.

The GP prior generalizes the covariate-free random effect. At the limit of large length-scales for all covariates, the squared-exponential term approaches a constant across all site pairs, so the prior reduces to the covariate-free random effect. Between these regimes, finite length-scales let covariate proximity govern how strongly sites are pooled.

#### 2.1.2 From calibrated scores to site-level vocal density

We define vocal density at site i, as the proportion of clips at that site containing vocalizations of the focal class. To estimate vocal density, we apply the site-level calibration to all unlabeled clips at site i, and combine the calibrated probabilities of unlabeled clips with any labeled clips from the site. Because inference is fully Bayesian, each posterior draw of the parameters yields a draw of vocal density, producing posteriors for vocal density at each site. We summarize these posteriors with highest posterior density credible intervals.

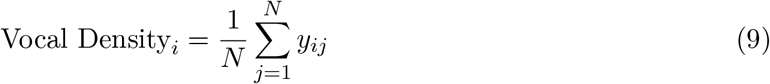

**Table 1.** Summary of notation used in the calibration model.

| Symbol | Definition |
| --- | --- |
| *Indices and data* |  |
| $i$ | Site index |
| $j$ | Clip index within a site |
| $N$ | Number of clips at a site |
| $S_{ij}$ | Logit-scale classifier score for clip $j$ at site $i$ |
| $y_{ij}$ | Binary clip label for clip $j$ at site $i$ |
| $\mathbf{x}_i$ | Vector of site-level covariates for site $i$ |
| *Calibration parameters* |  |
| $p_{ij}$ | Calibrated probability that clip $j$ at site $i$ contains the focal class |
| $\alpha_i$ | Site-level calibration intercept |
| $\beta_i$ | Site-level calibration slope (positive; $\beta_i = \text{softplus}(\tilde{\beta}_i)$) |
| $\tilde{\beta}_i$ | Unconstrained (pre-softplus) site-level slope |
| *Prior hyperparameters* |  |
| $\mu_\alpha, \mu_\beta$ | Prior means for the intercept and (pre-softplus) slope |
| $\sigma_\alpha, \sigma_\beta$ | Marginal standard deviations of the intercept and slope GPs |
| $\tau_\alpha, \tau_\beta$ | Per-site nugget standard deviations |
| $\ell_\alpha, \ell_\beta$ | Per-covariate GP length-scales controlling correlation decay |

#### 2.1.3 Implementation and inference

We fit the Gaussian-process calibration models using the No-U-Turn Sampler (NUTS), an adaptive variant of Hamiltonian Monte Carlo, implemented in NumPyro v0.20.0 (Python v3.13). We ran four independent chains in parallel, each with 1,000 warm-up and 1,000 sampling iterations, discarding the warm-up draws and retaining the 4,000 post-warm-up draws for inference. We placed the following hyperpriors on the GP parameters: *µ*_*α*_, *µ*_*β*_*∼ N* (0, 5) on the mean parameters, *σ*_*α*_, *σ*_*β*_*∼* HalfNormal(2) on the marginal standard deviations, and ***L***_*α*_, ***L***_*β*_*∼* Uniform(0.1, 10) on the per-covariate length-scales. To each GP covariance matrix we added an independent per-site nugget, *τ* ^2^**I** with *τ α, τ β ∼*HalfNormal(1), allowing sites with identical covariates to differ in calibration. Convergence was assessed using standard diagnostics, including trace-plot inspection, *R*^, and effective sample size. For each posterior draw we applied the fitted calibration to the unlabeled clips, yielding a posterior distribution of vocal density at each site.

#### 2.1.4 Performing ecological inference using site-level vocal densities

To quantify how site-level environmental covariates relate to vocal density, we used a two-stage cut procedure (Plummer, 2015). This propagates first-stage uncertainty into the regression while preventing the regression from feeding back to affect estimates of site-level vocal density. We deliberately avoid a joint specification: in a joint model the covariate–vocal density relationship being modeled would feed back to the first-stage site estimates. Any misspecification of that relationship would then propagate back into the site-level vocal densities. The first stage is the hierarchical Bayesian calibration model described above, which yields a posterior over site-level vocal densities. In the second stage, for each posterior draw we regressed logit(vocal density) on the site-level covariates by ordinary least squares (OLS), obtaining a coefficient estimate and standard error per covariate. We then sampled a coefficient value from its approximate within-draw posterior: the OLS estimate plus Gaussian noise with standard deviation equal to its standard error, and pooled these across draws. By the law of total variance, the resulting distribution combines between-draw variance (first-stage uncertainty in site-level density) and within-draw variance (second-stage regression error), yielding the cut posterior for each coefficient. We report the posterior mean, standard deviation, and 90% credible interval.

### 2.2 Data

We evaluate the method on three kinds of data:

- **Simulated data**, where the true vocal density is known exactly.
- **Two fully annotated field datasets**, where complete labeling allows us to know realized vocal density and lets us confirm our methods ability to estimate site-level vocal density from real classifier predictions.
- **A partially annotated field dataset**, which demonstrates the intended real-world application: ecological inference from a small number of labeled clips alongside extensive unlabeled audio.

#### 2.2.1 Simulated Data

We ran four simulated scenarios. For each scenario, we used a consistent dataset size of 100 sites, and 10,000 audio clips per site, representing a comparable size to a small-scale bioacoustic deployment. Classifier scores were generated from a mixture of Gaussian components with a shared standard deviation of 5: scores for negative clips (no vocalization) were drawn from a component with the lowest mean (*µ*_neg_ = *−*5), scores for true positives (vocalization present) from a component with the highest mean (*µ*_pos_ = 5), and, when a confusion class was present, scores for confusion clips from a component with an intermediate mean (*µ*_conf_ = 4), producing high-scoring false positives. We simulated two site-level covariates, each drawn independently from a standard normal distribution. In every scenario only the first covariate carries signal and the second is uninformative noise.

To reflect the fact that score distributions shift across sites, in all scenarios except the identical-sites baseline we jittered the component means on a per-site basis. Each site’s positive-and negative-component means were offset from the global values (*µ*_pos_, *µ*_neg_) by independent Gaussian noise with standard deviation 1. This introduces mild, unstructured variation between sites on top of the structured heterogeneity each scenario is designed to test. Full parameter values for all scenarios are given in the simulation code.

1. **Identical sites**. All sites share identical vocal density, and the same calibration. Scenario one represents an ecologically unrealistic scenario. This is the only scenario in which the assumptions underlying global calibration are met.
2. **Vocal density variation only, uncorrelated with covariates**. Sites vary in their vocal density, but not in any other factors (such as the presence of confusion sounds) that affect calibration. Variation in vocal density is not correlated with a covariate.
3. **Vocal density variation only, correlated with a covariate**. As in scenario two - sites vary in their vocal density, but here that variation is positively correlated with covariate one.
4. **Calibration variation due to a confusion class, correlated with a covariate**. All sites share the same vocal density, but the prevalence of a confusion class - which leads to
5. high-scoring ‘false positives’, varies. Confusion class prevalence is positively associated with covariate one.

#### 2.2.2 Fully annotated field datasets

For both fully annotated datasets, annotations were converted to window-level labels by dividing recordings into non-overlapping 5-second windows and treating a window as a positive for a species if an annotated vocalization of that species overlapped it by at least 0.25 seconds, or at least half of the vocalization’s total duration. Because annotation is exhaustive, the proportion of positive windows at a site is its realised vocal density, which we use as ground truth.

##### Pacific Northwest dawn chorus dataset

The Pacific Northwest dataset consists of 11.75 hours of fully annotated audio (n= 32,994 annotations) dawn chorus soundscape recordings from California, Oregon and Washington states in the Pacific Northwest, USA. 5 minutes of audio from 141 sites were annotated. The dataset was fully annotated for avian, mammalian and amphibian vocalizations, though we only use the 54 species of avian labels in our analysis. The data were collected with Wildlife Acoustics Song Meter SM4 ARUs (Wildlife Acoustics, Maynard, MA) programmed to record at 32kHz and 16-bit resolution during the hour following sunrise. The dataset also contains over 50 site-level covariates recorded at each site. For a full description of the covariates, see Weldy et al. (2024). We filtered these covariates to remove any with missing or infinite values, or that were discrete with fewer than 8 levels. Predictions were generated using the Google Perch bird vocalization classifier v1 (Google Research, 2023), which was trained on recordings from the Xeno-Canto community-science database.

##### Hawai’i dataset

The Hawai’i dataset consists of recordings collected at Hakalau Forest National Wildlife Refuge on the eastern slope of Mauna Kea, Hawai’i Island. These recordings were collected year-round at 10 sites along two elevational gradient transects using Songmeter SM4s (Wildlife Acoustics, Maynard, MA USA) programmed to record at 44.1kHz and 16-bit resolution. In addition to elevation, recording sites along the two transects vary in ecotype, however only elevation, which is known to be associated with vocal density of the target species (Navine et al., 2024a), was analysed as part of this study. Two species of Hawaiian honeycreeper were studied, the ‘Akiapōlā’au (*Hemignathus wilsoni*), and the ‘Alawī (*Loxops mana*), both of which are federally listed as endangered and considered endangered by the International Union for Conservation of Nature. We used predictions from a custom classifier with two classes for each species (adult and juvenile) trained on Google Perch classifier v1 embeddings for an earlier project.

#### 2.2.3 Partially annotated field dataset - Pennsylvania dataset

The Pennsylvania dataset consists of recordings collected at 283 sites across four public lands throughout the Central Appalachian region of Pennsylvania, USA. We investigated covariate effects on the vocal density of Wood Thrush (*Hylocichla mustelina*), which has experienced long-term population declines in eastern forests (Pardieck et al., 2020). Recordings were collected from a 10 day period of May 22 through May 31 (in either 2020 or 2021 depending on study site) using AudioMoth (v1.0/1.1) ARUs (Hill et al., 2019) programmed to record for two hours daily from 6–8 am EDT at a sampling rate of 32 kHz and medium gain setting. Predictions for the data were generated using the Google Perch classifier v1.

Labeled data for the Pennsylvania dataset were opportunistically drawn from a prior study conducted at the same sites (Lyon et al, *in review*). In that study, a convolutional neural network (CNN) classifier was trained using OpenSoundscape (Lapp et al., 2023), using strongly labeled audio clips from Xeno-Canto. Human listeners then labeled the highest-scoring clip per species per point per day. We use a total of 2 labeled clips from each site. The calibration set is therefore a non-random sample of the Perch score distribution - analogous to the logarithmic binning used in validation experiments.

Habitat covariates were measured at each point: basal area, woody stem density across two size classes, and leaf litter cover. Basal area (m^2^/acre) was estimated at plot center using a 10-factor cruising prism, counting all trees exceeding 10 cm diameter at breast height (DBH). Leaf litter cover was derived by averaging estimates from three 1 m^2^ quadrats - one placed near the plot center and one at the end of each of two transects extending 35 m outward. Transect bearings were drawn at random from three evenly spaced azimuths (0°, 120°, 240°). Woody stems below 10 cm DBH were tallied along both transects within paired 5 m^2^ segments, separated into short (< 1.5 m) and tall (> 1.5 m) categories.

### 2.3 Experiments

For the simulated scenarios and the fully annotated datasets, realized site-level vocal density is known. For these datasets we can use a subset of clips as labeled, estimate vocal density from the remaining (‘unlabeled’) clips, and compare the estimates against truth. The Pennsylvania dataset has no ground-truth for vocal densities and is used instead to demonstrate ecological inference in a realistic, sparsely labeled setting.

#### 2.3.1 Held-out labeling experiments

In each experiment, clips within a labeled site were selected for labeling using logarithmic binning of the observed score quantiles (Navine et al., 2024b), which concentrates labeling effort on high-scoring clips. Each selected clip received a 0/1 label for presence of the focal species. For each labeled calibration set we fit our hierarchical model and both baseline methods (see below), then aggregated the calibrated per-clip probabilities together with known labels from the calibration set to obtain a posterior distribution of vocal density on the unlabeled clips at every site.

##### Simulated scenarios

For each of the four scenarios we repeated the full simulation procedure across 10 replicate datasets. In each replicate we randomly assigned half of the 100 sites to the calibration set and reserved the other half as test sites with no labeled data. This lets us verify that interval coverage extends to sites without labeled data. We applied one fixed level of labeling effort: a total labeling budget of 250 clips, selected by logarithmic binning. Each scenario therefore yields 100 sites × 10 replicates = 1,000 site-level estimates of vocal density, over which we report averaged evaluation metrics.

##### Fully annotated field datasets

For the Pacific Northwest and Hawai’i datasets we instead varied labeling effort, repeating the procedure across a range of labeling effort. At each level, for each species, we generated 3 random calibration sets. Evaluation metrics are averaged across all species. We report each metric as a function of labeling effort, showing how point-estimate accuracy, interval precision, and coverage change as labeling effort changes.

#### 2.3.2 Baseline calibration methods

We compared our hierarchical model with two standard alternatives, both fit in the same Bayesian framework (NumPyro) with weakly informative priors: global Platt scaling, with a single intercept and slope shared across all sites; and independent site-level Platt scaling, with a separate intercept and slope per site and no pooling. For all calibration models we used a Normal(0, 5) prior for the intercept and a Half-Normal(2) prior for the slope. We followed the same procedure as described in our hierarchical modeling procedure: fitting the calibration model, aggregating the calibrated per-clip probabilities on unlabeled data (plus the known labels for labeled data) to yield a posterior over site-level vocal densities. Global Platt scaling assumes that calibration is identical at all sites. Independent site-level Platt scaling enforces the opposite assumption of no information sharing. Our hierarchical model interpolates between these two.

#### 2.3.3 Evaluation metrics

Each method for estimating site-level vocal density on the simulated and fully annotated audio datasets is evaluated using three metrics: mean squared error (MSE), credible interval (CI) width, and observed coverage of site-level ground truth. MSE was used to quantify overall accuracy of point estimates for site-level vocal density. CI width was used to assess the precision of each method, with narrower intervals indicating more precise estimates. Observed coverage was defined as the proportion of site estimates in which the CI contained the true parameter value. Together, these metrics provide a comprehensive assessment of both point estimation accuracy and interval performance across calibration approaches. Because all three metrics require known truth, they apply only to the simulated and fully annotated datasets.

#### 2.3.4 Demonstration: ecological inference on partially annotated data

To illustrate the method in its intended setting, we applied it to the Pennsylvania dataset, using two labeled clips per site to calibrate the Perch scores and estimate Wood Thrush vocal density at all 283 sites. We then applied the second stage regression (see Model section “Ecological Inference”) to relate vocal density to the four habitat covariates. No ground truth vocal density is available here, so this analysis demonstrates ecological inference rather than validating accuracy. We expect to find that Wood Thrush vocal density is positively associated with structural features of mature mesic deciduous forest, such as basal area, leaf-litter cover and tall stem density (Bertin, 1977; Evans et al., 2020).

## 3 Results

### 3.1 Simulation Experiments

The central result of our simulations is that modelling between-site variation lets the hierarchical model recover accurate, well-calibrated vocal density estimates at a fraction of the labeling effort demanded by site-level calibration. Importantly, it does so even at sites with no labeled data at all (Figure 2, 3). Across all four simulated scenarios (Figure 2), our proposed hierarchical model achieves close to the desired 90% credible interval coverage (observed coverage across all simulations 91.8% when fit without covariates, 94.5% with covariates) while maintaining lower MSE and narrower CIs than site-level calibration. Relative to site-level calibration at the same labeling effort, the hierarchical model (with covariates) reduced MSE by 76.4–97.9% and narrowed CIs by 63.5–68.2% across the four scenarios. In scenarios where site-level heterogeneity was driven by covariates (scenarios three and four), the use of covariates improved performance: MSE fell by 58.8% (scenario 3) and 57.9% (scenario 4) and CIs narrowed by 19.1% and 16.5%, respectively, relative to the hierarchical model without covariates.

**Figure 2.**
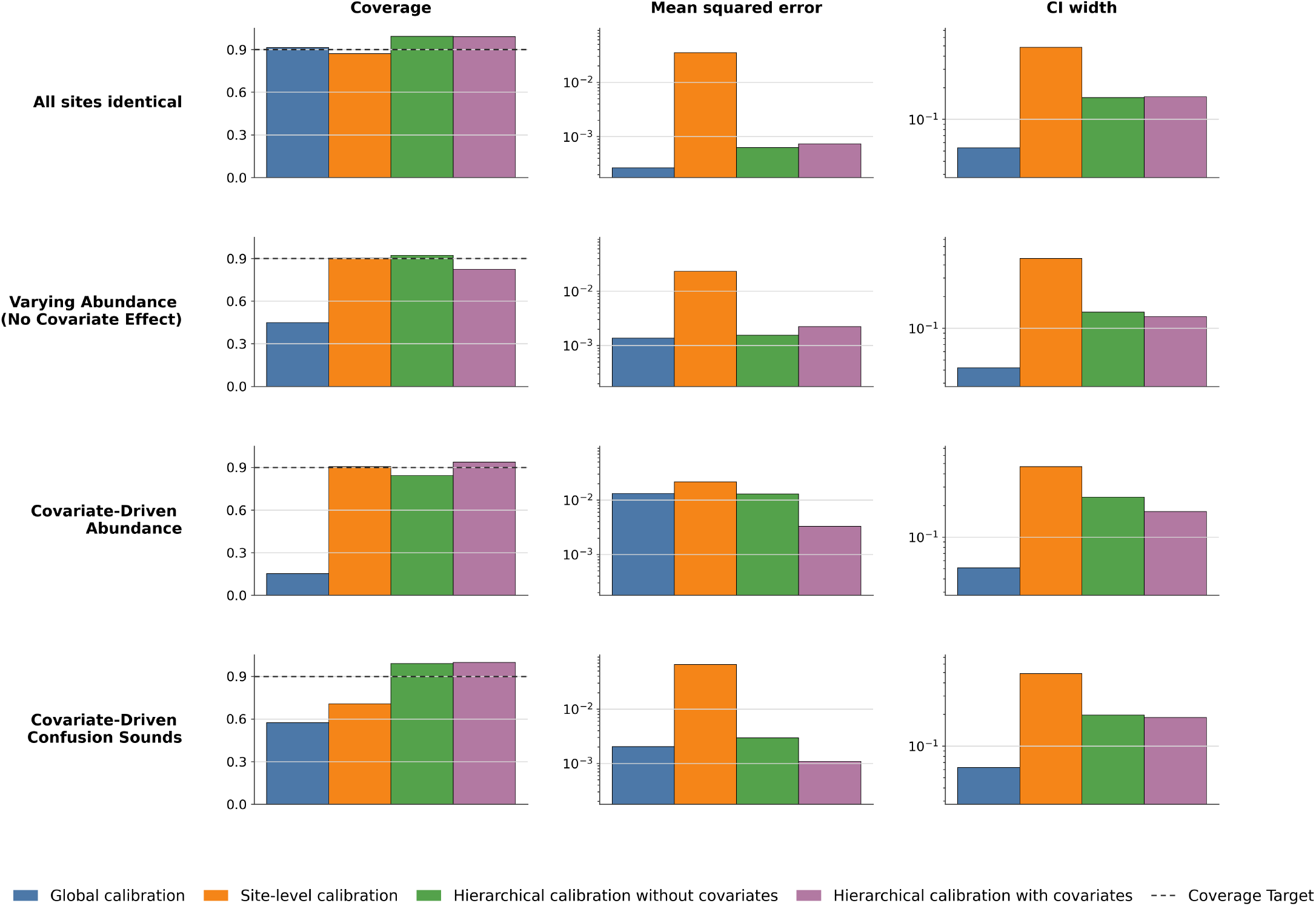
Simulation study comparing calibration methods. Each row represents a different data-generating simulation: (1) all sites identical with no variation in abundance, (2) site-varying abundance independent of covariates, (3) covariate-driven variation in abundance between sites, and (4) covariate-driven variation in confusion sounds between sites. Columns show three evaluation metrics: 90% credible interval coverage (left), mean squared error (center), and CI width (right). The dashed line in coverage plots indicates the target of 90% coverage. MSE and CI width are displayed on log scales. Results are averaged across all 10 simulation runs. The hierarchical model with covariates achieves close to the targeted 90% coverage across all scenarios while maintaining low MSE and narrow credible intervals, particularly when covariate effects drive site-level heterogeneity.

**Figure 3.**
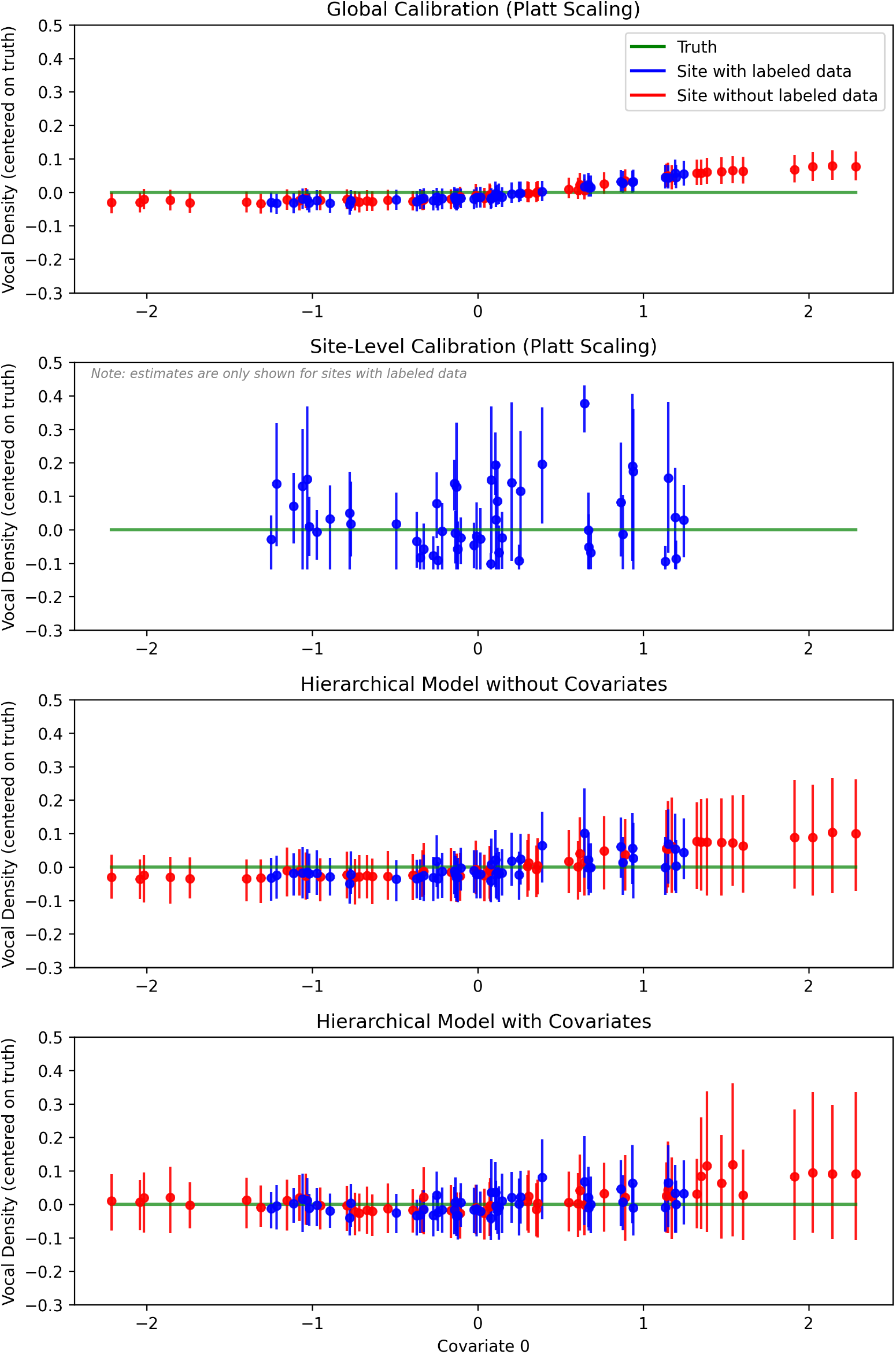
Model performance under covariate-driven variation in confusion sounds. Vocal density estimates for each site in simulated scenario four - where the prevalence of a confusion sound varies across sites, increasing with Covariate 1. Estimates are centered on the true value (green line at zero). Blue points indicate sites with labeled validation data; red points indicate sites without labeled data. Error bars represent 90% credible intervals. Global calibration applies a single calibration across all sites and cannot account for site-level variation in false positive rates, leading to CIs that do not attain coverage. Site-level calibration can only be used for sites that were included in the calibration set. The hierarchical model without covariates partially pools information across sites but does not use the information in covariates. In contrast the hierarchical model with covariates captures the covariate-driven pattern, yielding estimates closer to truth and narrower credible intervals, particularly at sites 1w4ithout labeled data.

The only simulated scenario in which global calibration attained desired coverage is simulation one – where all sites are simulated to be identical. In scenarios 2, 3 and 4 where heterogeneity between sites was simulated, observed coverage of global calibration was 42.0%, 15.3% and 52.4% respectively. Site-level calibration achieved coverage between 0.745 and 0.921 across the four scenarios, but produced much wider CIs (0.459–0.497) and higher MSE than the hierarchical model at the same labeling effort.

In scenarios three and four, where site-level heterogeneity is driven by covariates, incorporating covariates into the hierarchical model further reduces MSE and narrows CIs relative to the hierarchical model without covariates. In these scenarios, the hierarchical model without covariates still attains close to nominal coverage, but the covariate-informed model produces more precise estimates with narrower CIs. In scenarios one and two, the covariates are uninformative. The model with and without covariates produce similar estimates: coverage was comparable (0.991 vs. 0.993 in scenario 1; 0.933 vs. 0.931 in scenario 2), and differences in MSE and CI width were small.

The advantage of our model is starkest at sites with no labeled data (Figure 3). Site-level calibration has no labeled data to fit at such sites, so it cannot produce an informed estimate; global calibration can, but its credible intervals failed to attain coverage. Only the hierarchical model maintained close to nominal coverage there, because it alone carries information about between-site variation learned from labeled sites across to unlabeled ones.

### 3.2 Fully Annotated Datasets

The learned calibration curves for different classes vary in their sensitivity to covariates (Figure 4). For example, in the Hawai’i dataset for adult ‘Akiapōlā’au and juvenile ‘Alawī (Figure 4 panels A, D), calibration curves were relatively consistent across sites. While calibration curves for juvenile ‘Akiapōlā’au and adult ‘Alawī (Figure 4 panels B, C) varied substantially across sites along the elevational gradient.

**Figure 4.**
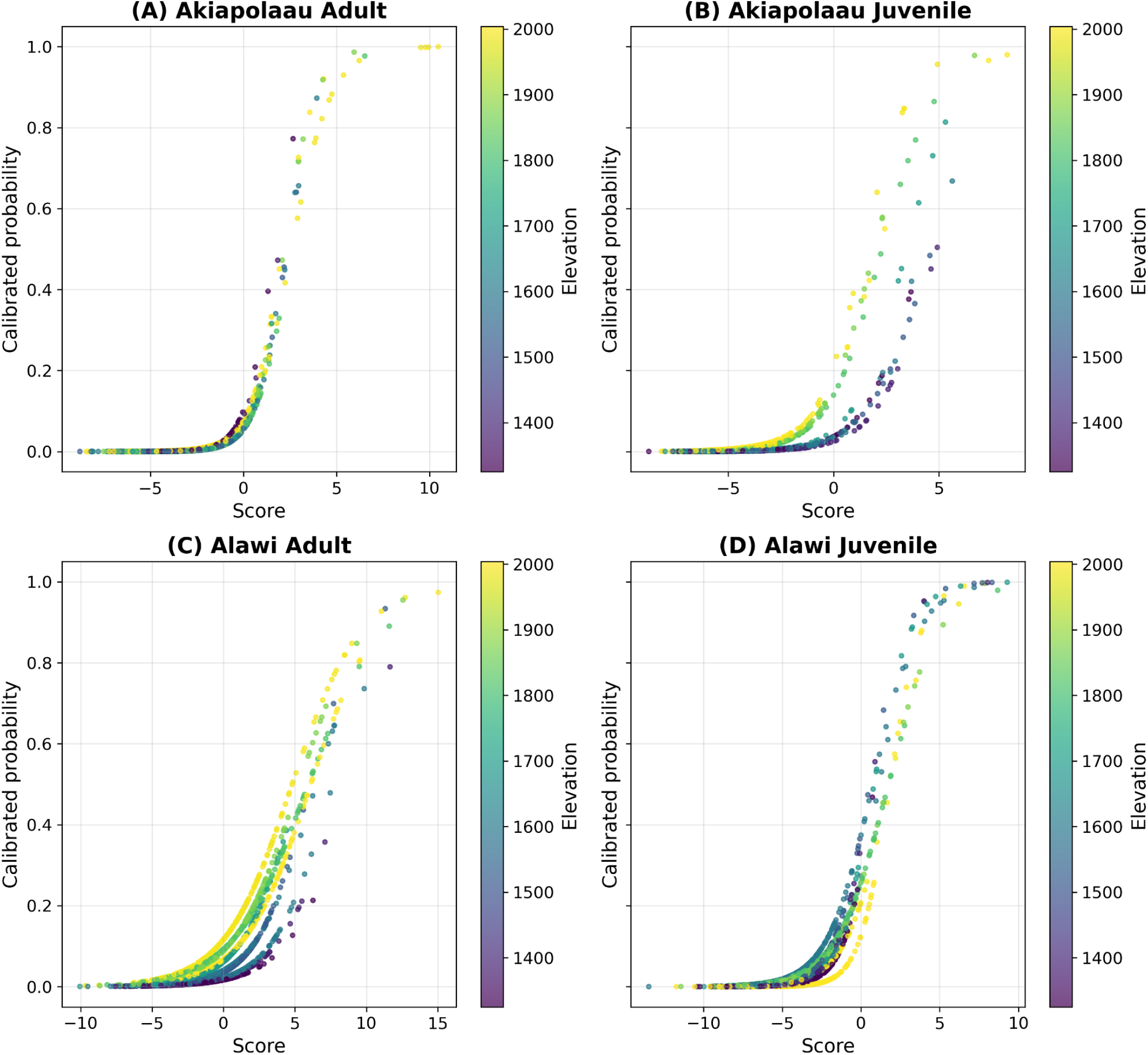
Calibrated probabilities for two species in two life stages across sites on an elevation gradient. We can see in (A) and (D) little variation on the learnt calibration curve between sites, indicating adult ‘Akiapōlā’au and juvenile ‘Alawī calibration is not heavily influenced by elevation. However, for panels (B) and (C) calibration curves change more significantly between sites, revealing elevation’s influence on calibration for juvenile ‘Akiapōlā’au and adult ‘Alawī.

In the field data, across the Hawai’i and Pacific Northwest field datasets (Figure 5), MSE and CI width decreases with increased labeling effort for all calibration approaches. In the Hawai’i dataset, global calibration provides the lowest MSE estimates when the number of labeled datapoints is below 100, whilst in the Pacific Northwest dataset our hierarchical model provides the lowest MSE at low levels of labeling effort. Site-level calibration and our hierarchical model achieve close to desired coverage. The hierarchical model achieves coverage with narrower CIs than site-level calibration.

**Figure 5.**
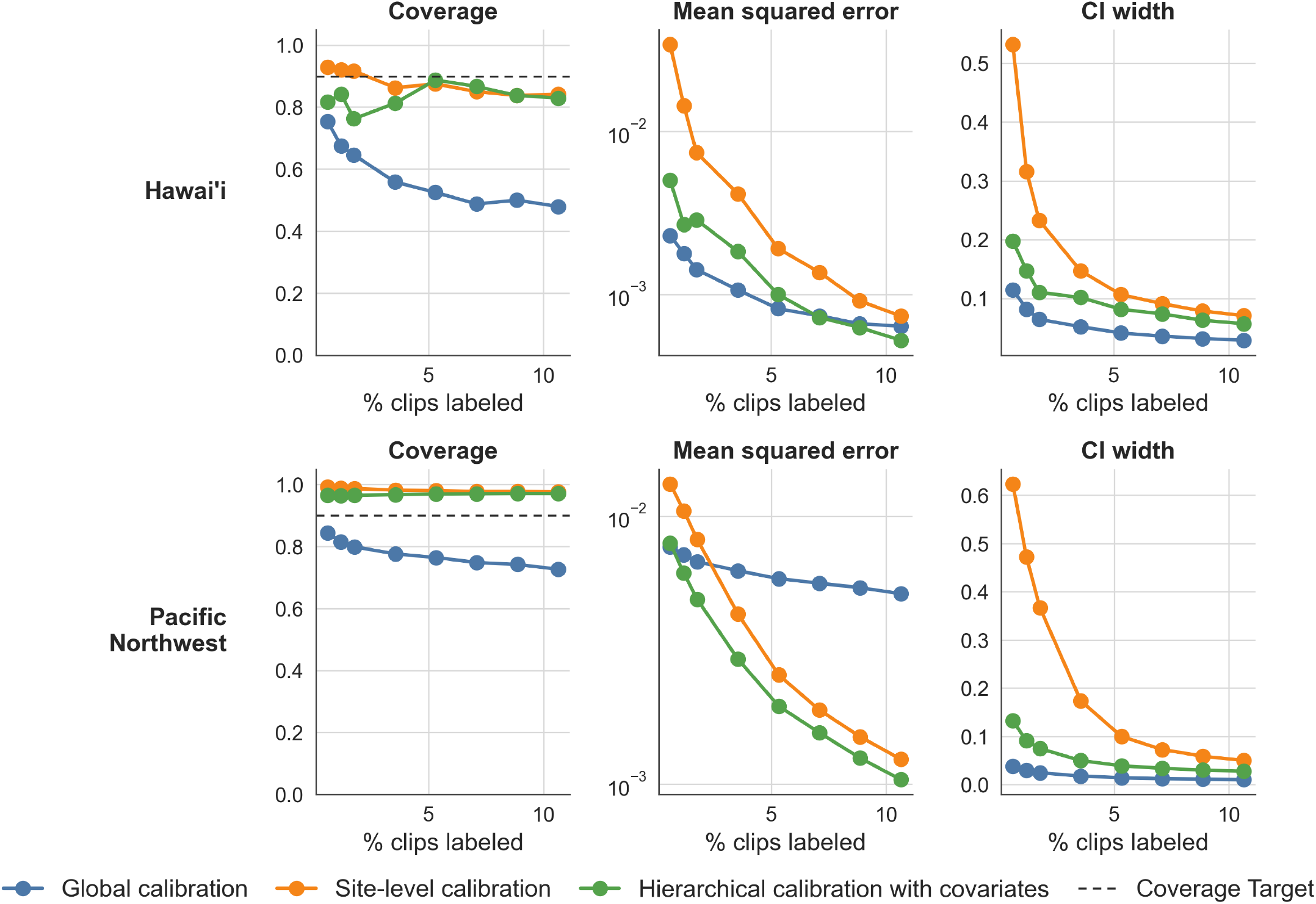
Performance of our calibration method averaged over each class in the Hawai’i and Pacific Northwest datasets. Each column shows a different evaluation metric: mean squared error (MSE) of predicted site-level vocal densities, average width of the 90% credible intervals and observed coverage of the 90% posterior credible intervals. The x axis shows the percentage of clips in the dataset used as labeled data points. For each level of labeling effort, data were randomly sampled for labeling using log-binning, with 3 different random seeds.

### 3.3 Partially annotated field dataset

Applying the two-stage approach for ecological inference, we estimated covariate effects on vocal density at the 283 Pennsylvania sites for Wood Thrush. Basal area (posterior mean *β* = 0.84, 90% CI [0.42, 1.25]) and the density of tall woody stems (*β* = 0.85, 90% CI [0.45, 1.24]) were positively associated with vocal density, with credible intervals excluding zero. Short woody-stem density (*β* = 0.29, 90% CI [*−* 0.14, 0.71]) and leaf litter cover (*β* = 0.09, 90% CI [*−*0.34, 0.49]) had credible intervals overlapping zero. Effects are reported on the logit scale; full posterior distributions are shown in (Figure 6).

**Figure 6.**
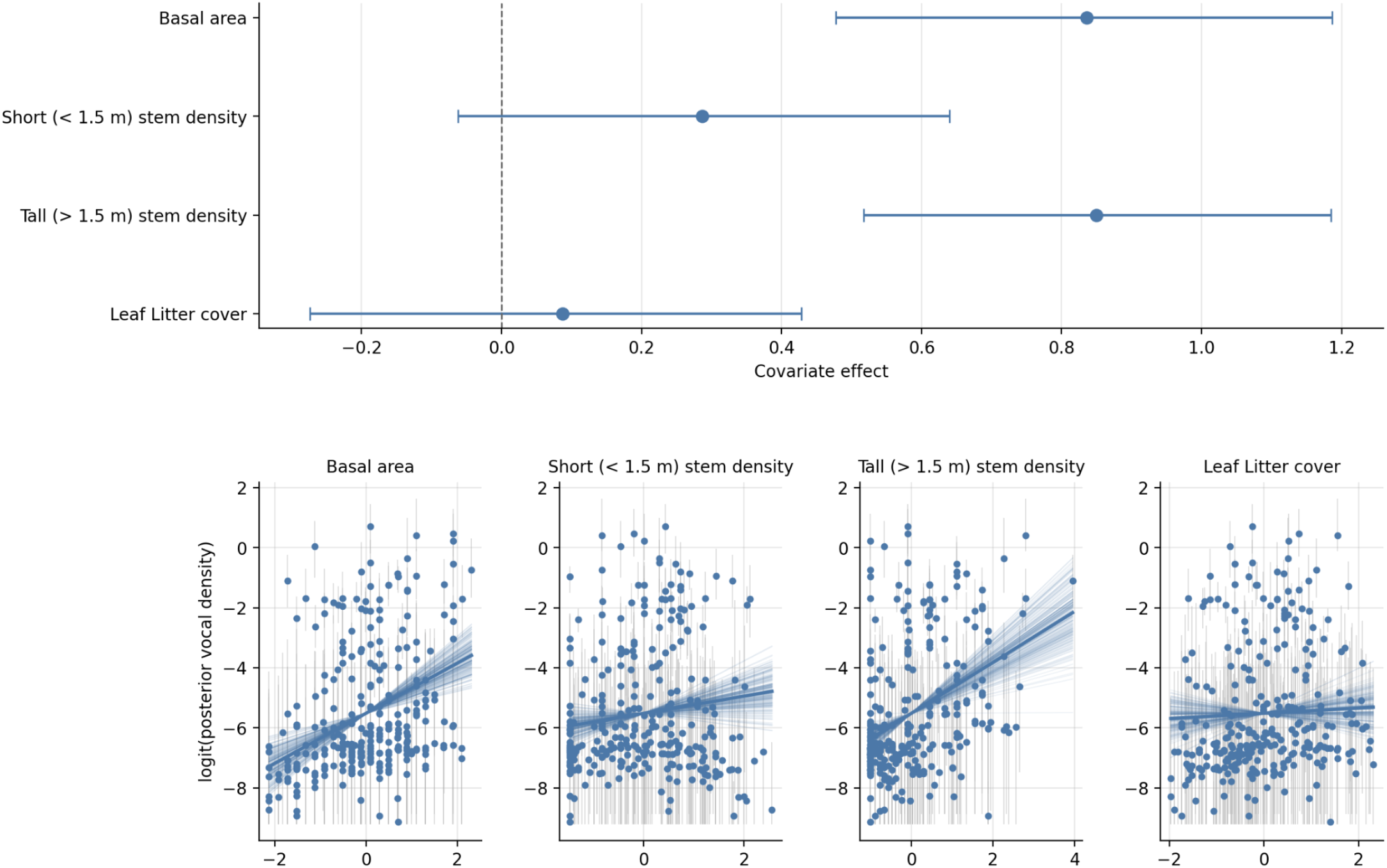
Covariate effects on Wood Thrush vocal density estimated by two-stage inference. (A) Forest plot of stage-2 multivariate regression slopes. Points and bars show the posterior mean and 90% credible intervals; the dashed vertical line marks zero. (B) For each covariate, posterior mean logit(vocal density) per site (grey bars: 90% credible interval) plotted against the z-scored covariate value. The bold blue line shows the posterior-mean partial slope; faint lines show 200 draws from the joint posterior, illustrating slope uncertainty. All panels report the results of a single joint multiple regression.

## 4 Discussion

Our hierarchical calibration model yields site-level vocal density estimates while reducing the data labeling burden compared to fitting separate site-level calibrations. Importantly, it quantifies the uncertainty in these estimates with CIs that attain coverage, even when vocal density is estimated at sites from which no data has been labeled. On a real dataset, it recovers established ecological relationships from as little as two labeled clips per site. As passive acoustic datasets grow increasingly large (Darras et al., 2025) and expert labeling time remains limited, producing calibrated site-level estimates in a label-efficient manner is increasingly important.

Both our simulation and field results demonstrate the pitfalls of performing global calibration and then estimating site-level vocal density. In most ecological studies, sites are the natural subgroupings ecologists care about, however this pattern of subunits exhibiting “internal domain shift” relative to the dataset as a whole is related to the problem of “multicalibration” in the machine learning literature (Hebert-Johnson et al., 2018) - the requirement that a classifier’s scores be calibrated not just on average, but within identifiable subgroups. Global calibration assumes no domain shift between sites and yields CIs for site-level estimates that are overly narrow. The effect on the point estimate is variable: in the Hawai’i dataset the bias introduced by global calibration lowered MSE at the lowest annotation effort, whereas in the Pacific Northwest dataset it raised MSE, but in both cases the CIs fail to appropriately estimate uncertainty. The same issue is known to affect approaches that choose a fixed threshold across all sites (Metcalf et al., 2022; Knight et al., 2017). Independent site-level calibration does attain desired coverage given sufficient labeled data, but is limited in two important respects: at equal labeling effort it produces wider CIs than our hierarchical model, because it cannot borrow strength across sites; and labeling effort scales linearly with the number of sites - it can only produce estimates for sites with labeled data. In Figure 3, for example, our hierarchical model produces estimates for all sites, whereas site-level calibration cannot be used at sites without labeled data which, in a large deployment with a limited labeling budget, will be the majority.

Our work is related to false-positive models (e.g., Clement et al., 2022; Doser et al., 2021; Udell et al., 2024). Our approach differs, however, in that it exploits the continuous score outputs of classifiers rather than collapsing them to a thresholded binary detection/non-detection. In this respect it resembles recently proposed continuous-score occupancy models (Rhinehart et al., 2022; Kéry and Royle, 2021). The modeling approach we take, however, is discriminative: we model the posterior class probability *P* (*y* = 1|*s*) as a function of classifier score and estimate vocal density from these calibrated probabilities. The proposed continuous-score occupancy models instead take a generative approach, fitting class-conditional score densities *p*(*s* | *y* = 1) and *p*(*s* | *y* = 0) as Gaussians. Under a generative modeling approach, vocal density would be estimated from the mixing proportion of the two distributions. The two approaches differ importantly in the strength of their assumptions: estimates from the generative approach depend on correctly specified class-conditional densities, and if either is misspecified, the estimate may be biased and its CIs may not attain desired coverage. In contrast the discriminative approach makes weaker assumptions, requiring only that *P* (*y* = 1|*s*) be adequately approximated by a sigmoid. This robustness comes at a cost in efficiency: when the class-conditional densities are correctly specified, fitting the full likelihood is more efficient (Efron, 1975); when they are not, logistic regression is the more reliable choice (Press and Wilson, 1978). We prioritise robustness of coverage. Our simulations demonstrate this robustness: in scenario 4, the presence of a high-scoring confusion sound component means the sigmoid is misspecified, yet the CIs still attain desired coverage.

Care should be taken when performing ecological inference using site-level vocal densities produced by our method. Our site-level estimates are posterior distributions. Regressing point estimates against covariates of interest would discard that uncertainty and, by treating dependent estimates as independent observations, amounts to pseudoreplication. Downstream inference requires propagating uncertainty into the regression step. In our Pennsylvania case study we achieve this with a two-stage approach rather than a single joint model. We use this cut-style approach because, in a joint specification, the parametric form assumed for the relationship between covariates and vocal density would feed back into site-level estimates; keeping the two stages separate prevents this feedback. The cost of performing a separate second-stage regression is that the hierarchical model in the first stage shrinks site-level estimates toward the global mean, biasing estimated covariate effects in the second stage regression toward zero. The magnitude of shrinkage depends on the precision of site-level estimates, which improves with greater labeling effort and the skill of the classifier. Shrinkage of effect sizes is therefore most pronounced when labels are scarce and the classifier is weak. This effect works against detecting an association rather than creating a spurious one.

The covariate effects inferred from our Pennsylvania dataset recapitulate known habitat associations for Wood Thrush. Basal area and the density of tall (>1.5 m) woody stems both had positive associations with vocal density and CIs that excluded zero, consistent with the species’ preference for mature mesic deciduous forest with a well-developed subcanopy and shrub layer (Bertin, 1977; Evans et al., 2020). That we recovered these established relationships from only two labeled clips per point - despite shrinkage being greatest at low labeling effort as described above - demonstrates that our approach retains power to detect real associations even under aggressive label efficiency, and illustrates the kind of inference at scale our model makes tractable.

### Using our method

1. **Choose covariates**. Identify site-level covariates that you wish to do inference on, or that you believe drive between-site variation in classifier calibration. Including covariates that carry no signal was not harmful in our simulation experiments, because the Gaussian process down-weights them through its per-covariate length-scales, so it is reasonable to favour including plausible candidates.
2. **Label the calibration set**. The exact amount of labeling effort will depend on the desired precision of vocal density estimates, classifier quality and dataset size. We suggest labeling at least 200 clips per class. Clips should be selected from multiple sites. We suggest using logarithmic binning of the score distribution (Navine et al., 2024b), to select 5 clips per-site, from at least 40 sites. Site selection can be stratified across covariate space. Consider also other uses of the labeled data: for example for maximal information about occupancy, you may wish to label the top-scoring clip from each site, in addition to the stratified sample.
3. **Fit the model and estimate vocal densities**. Both the labeled (score, label) and unlabeled data are passed to the calibration model to obtain a posterior distribution over vocal density at every site, including sites with no labeled data. Confirm convergence of vocal density estimates using standard diagnostics such as *R*^, effective sample size, and trace-plot inspection.
4. **Perform second-stage inference (optional)**. If the goal is to relate vocal density as a proxy for abundance to environmental covariates, propagate first-stage uncertainty using the two-stage cut procedure rather than regressing point estimates, which would discard uncertainty. Bear in mind that shrinkage of estimates biases the estimated effects toward zero, most strongly when labels are scarce or the classifier is weakest.

There are several limitations to our approach. First, though motivated by correcting for between-site variation in classifier performance driven by covariates, we cannot correct for every source of covariate-associated variation in the sampling process. If, for example, proximity to a stream is associated with background noise that reduces a recorder’s effective detection radius, this is a sampling bias that cannot be addressed with our calibration approach. Second, the advantage of borrowing strength is greatest when site-level heterogeneity is at least partly explained by the available covariates; we expect the largest gains where calibration varies across sites in ways covariates capture. Third, although large-scale deployments are what motivate label-efficient calibration, the computational cost of fitting a Gaussian process scales cubically with the number of sites (Golding and Purse, 2016). Our largest application here comprises only 283 sites. Computation at the largest deployments will likely require sparse or inducing-point approximations, variational inference, or other methods for reducing computational intensity (Ingram et al., 2020).

There are a number of important future directions related to our work. How best to spend a limited labeling budget remains an open question. Throughout, we selected clips using the logarithmic binning of Navine et al. (2024b) which heuristically concentrates effort on high-scoring clips likely to be positives. A principled future direction would embed the labeling within an active-learning framework (Settles, 2012), using the hierarchical model itself to identify the labels that would most reduce uncertainty in site-level vocal densities or in coefficients of ecological interest. This is a rich area for future work, as identifying which clips to label requires balancing each datapoint’s utility: for calibration of existing models, for training of future models and for the ecological information resolving its label carries in its own right (Kurinchi-Vendhan and Beery, 2026). There is a perceived trade-off between using labels to calibrate an existing model and using them to train or fine-tune a new one. We argue this trade-off is less stark than it appears: vocal density is a property of the data rather than the model, so evidence about it can accumulate across successive models. The posterior over vocal density from one model encodes information contained in the labels in the calibration set, and can serve as a prior for future analyses - even if those same labels are used to train the next iteration of the classifier. The one firm constraint is that no label may be used to both train and calibrate the same classifier, since a model’s scores on its training data are over-optimistic. Methods that integrate information from labeled data and predictions from multiple models on unlabeled data (Shanmugam et al., 2025) are an interesting route forward.

An obvious extension is a community model that calibrates many classes jointly rather than independently. Adding a hierarchical component over species - treating species-level calibration parameters as draws from a shared distribution - would let the model borrow strength across species, improving estimates across classes. Such a model could also use the full vector of classifier scores, including non-focal classes, as predictors: scores for acoustically similar classes carry information about likely confusion errors, and could improve calibration where such confusions are prevalent.

The pairing of autonomous sensors with machine-learning classifiers is widely expected to expand biodiversity monitoring to spatial and temporal scales unattainable by manual survey (Besson et al., 2022; Tuia et al., 2022; Pollock et al., 2025). Realising that promise at scale depends on turning raw classifier outputs into ecologically meaningful quantities with accurately quantified uncertainty — and that uncertainty matters in its own right, since estimates that ignore it can leave managers unable to weigh decision risks or even to detect the population changes their monitoring is meant to capture (Milner-Gulland and Shea, 2017). The hierarchical calibration framework we present addresses this bottleneck directly: it converts classifier scores into calibrated, site-level estimates of vocal density, quantifies their uncertainty with intervals that attain coverage, and does so with less required human expert labeling effort.

## 5 Acknowledgements

We thank Tessa Rhinehart, Sam Lapp, Lauren Chronister, Cameron J Fiss, Chapin Czarnecki, Leonardo Viotti, Patrick Lyon and Yiluan Song for comments on drafts of this manuscript. We thank Patrick Lyon, Chapin Czarnecki and Nickolus Stahlman for the labeling of data from the Pennsylvania dataset. We would like to thank the Pennsylvania Game Commission and the Pennsylvania Department of Conservation and Natural Resources for enabling land access and logistical support for collection of the Wood Thrush data. This material is based upon work supported by the National Fish and Wildlife Foundation. This research was also supported in part by the University of Pittsburgh Center for Research Computing through the resources provided. LLMs (multiple models) were used for writing code for conceptual Figure 1 and for some of the analysis.

## 6 Author contributions

All authors made substantial contributions to conceptualization and design. LFH and TD contributed code development and testing. AKN, MW & TD contributed to acquisition of data. JK acquired funding. All authors contributed to conceptualization, analysis and interpretation of data, and drafting of the article. All authors contributed critically to the drafts and gave final approval for publication.

